# Postoperative Stress Accelerates Atherosclerosis through Inflammatory Remodeling of the HDL Proteome and Impaired Reverse Cholesterol Transport

**DOI:** 10.1101/2025.05.02.651357

**Authors:** Dominique M. Boucher, Valerie Rochon, Thomas Laval, Victoria Lorant, Abigail Carter, Christina Emerton, Nathan Joyce, Nysa Vinayak, Marlena Scaffidi, Rebecca C. Auer, Scott M. Gordon, Mireille I. Ouimet

**Affiliations:** Department of Biochemistry, Microbiology and Immunology, University of Ottawa, 451 Smyth Rd., Ottawa, ON, K1H 8M5, Canada; University of Ottawa Heart Institute, 40 Ruskin St, Ottawa, ON, K1Y 4W7, Canada; Saha Cardiovascular Research Center and Department of Physiology, University of Kentucky, 741 South Limestone, Lexington, KY, 40536-0509, USA; Cancer Therapeutics Program, Ottawa Hospital Research Institute, 501 Smyth Rd, Ottawa, ON, K1H 8L6, Canada; Division of General Surgery, Department of Surgery, University of Ottawa, 501 Smyth Rd, Ottawa, ON, K1H 8L6, Canada

**Keywords:** surgery, HDL, atherosclerosis, reverse cholesterol transport, acute phase response

## Abstract

**BACKGROUND:** Over 10 million patients undergoing non-cardiac surgery annually experience major cardiovascular complications within 30 days, many due to destabilized atherosclerotic plaques. Reverse cholesterol transport (RCT), a key pathway for cholesterol removal by HDL and apoA-I, is critical in preventing plaque progression. While surgery-induced inflammation is known to impair HDL function, its effects on RCT and plaque stability remain unclear.

**METHODS:** To isolate the impact of surgical inflammation, independent of blood loss, we developed an abdominal laparotomy model in *apoE*^*-/-*^ mice on a Western diet, minimizing blood loss and avoiding perioperative blood sampling. We assessed plasma cholesterol efflux capacity, performed proteomic analysis of HDL, and analyzed atherosclerotic plaques for lipid content, perilipin-2 (PLIN2), cleaved-caspase-3 (c-Casp-3), and necrotic core expansion. A novel dual-label, dual-cell-type *in vivo* RCT model was developed to compare RCT from macrophage-derived (BMDMs) and vascular smooth muscle cells (VSMCs)-derived foam cells. Recombinant apoA-I (rApoA-I) was tested for therapeutic rescue of impaired RCT.

**RESULTS:** Surgery significantly reduced RCT for at least 48 hours, paralleled by a drop in cholesterol efflux capacity and inflammatory remodeling of HDL, marked by elevated serum amyloid A (SAA1/2) and reduced apoA-I. Plaques showed a 1.6-fold increase in intracellular lipids and PLIN2 expression at 24 hours post-surgery, with elevated c-Casp-3 indicating lipid-driven apoptosis. Foam cell analysis revealed increased PLIN2 in both CD45^+^ (leukocyte) and CD45^-^ (non-leukocyte) subtypes, with leukocyte foam cells expressing higher PLIN2. c-Casp-3^+^ apoptotic cells were predominantly PLIN2^high^ and of both leukocytic and non-leukocytic origin. By day 15, the necrotic core area increased by 1.5-fold with sustained loss of plaque cellularity. Using our dual-cell-type RCT model, we found that surgery significantly impaired BMDM RCT *in vivo*, while VSMC RCT remained largely unaffected, highlighting foam cell subtype-specific vulnerability to surgical inflammation. These findings were mirrored in general surgery patients, whose postoperative plasma exhibited markedly reduced cholesterol efflux capacity. In mice, rApoA-I treatment partially restored RCT and reduced plaque lipid accumulation.

**CONCLUSIONS:** Surgical inflammation acutely impairs HDL function and RCT, triggering lipid accumulation, foam cell apoptosis, and accelerated plaque destabilization independent of blood loss. Immediate restoration of apoA-I at the time of surgery, aiming to counteract the acute phase response, may offer a targeted strategy to reduce postoperative cardiovascular risk.

## INTRODUCTION

Each year, over 10 million patients undergoing non-cardiac surgery worldwide will suffer major cardiovascular complications within 30 days of their procedure^1,2^. Even though atherosclerosis, affecting 30% of all surgical patients, is a known risk factor for major postoperative cardiovascular complications and mortality^1-4^, the triggers for plaque disruption and rupture during the perioperative period remain unclear. Atherosclerosis is a chronic sterile inflammatory disease characterized by the build-up of lipid-rich plaques in the arteries. Initially, the infiltration and modification of low-density lipoprotein (LDL) in the vascular wall elicits the recruitment of circulating monocytes that differentiate into macrophages and engulf modified LDL, transforming into foam cells^5-7^. In parallel, a subset of neighboring vascular smooth muscle cells (VSMCs) undergoes lipid loading and phenotype switching to macrophage-like foam cells^8-13^. The recent discovery that VSMC foam cells can make up over 70% of the foam cell population in both mouse and human atherosclerotic plaques^9-13^ challenges the previous view that macrophages are the primary source of foam cells and underscores the need to investigate VSMC foam cells in atherosclerosis.

In atherosclerosis, the balance of pro-inflammatory and inflammation-resolving mechanisms dictates the final clinical outcome^14^. Cholesterol efflux pathways exert antiatherogenic and anti-inflammatory effects by reducing foam cell cholesterol accumulation and by limiting activation of the NLRP3 inflammasome^15,16^. In parallel, efferocytosis, the clearance of apoptotic cells by macrophages, helps prevent secondary necrosis and stimulates the release of anti-inflammatory cytokines^17^. In advanced atherosclerosis, inflammation-resolving mechanisms become overwhelmed and uncleared apoptotic cells become secondarily necrotic, promoting inflammation and tissue damage. Additionally, macrophage-VSMC crosstalk dictates plaque stability^18^. While pro-resolving factors released by macrophages in early atherosclerosis promote VSMC proliferation to increase plaque stability^17^, pro-inflammatory mediators released by macrophages in late atherosclerosis cause VSMC de-differentiation, extracellular matrix degradation and fibrous cap thinning^18^. Ultimately, impaired inflammation resolution and expansion of the necrotic core leads to plaque disruption, acute thrombosis, and tissue ischemia or infarction^19^.

The 1948 Framingham Heart Study was the first to report an inverse association between plasma high density lipoprotein cholesterol (HDL-C) and cardiovascular disease (CVD) risk^20^. This discovery spurred numerous studies into the mechanisms by which HDL protects against CVD, leading to the identification of the reverse cholesterol transport (RCT) pathway^21^. RCT, the net flux of cholesterol from peripheral tissues, including arterial foam cells, to the bloodstream for fecal excretion, is crucial for protecting against atherosclerosis^22^. Together, the ATP-binding cassette (ABC) transporters ABCA1 and ABCG1 mediate the bulk of cholesterol efflux to HDL, with ABCA1 transferring cholesterol to apolipoprotein A-I (apoA-I) and small HDL particles, and ABCG1 facilitating efflux to mature HDL^23-28^. While negative HDL-raising clinical trials have cast doubt on the ‘HDL hypothesis’, it has become clear that plasma HDL-C levels per se are not a faithful biomarker of CVD and do not accurately reflect HDL particle abundance, the distribution of HDL subspecies, or the ability of HDL to mediate RCT^22^. Indeed, large clinical studies showed that HDL cholesterol efflux capacity (CEC) has a stronger inverse relationship with CVD risk than HDL-C levels, even after adjusting for HDL-C^29-32^.

Strong evidence supports HDL’s anti-inflammatory role beyond RCT, contributing to its anti-atherosclerotic effects^33-35^. Meta-analysis of HDL proteomes from 45 studies identified many inflammation-related proteins, including acute phase response (APR) proteins serum amyloid A (SAA) 1 and 2, indicating HDL’s broader functions^33^. Acute inflammation can induce changes in HDL composition and metabolism^36^, impairing RCT^37,38^. Mechanistically, acute inflammation markedly raises HDL SAA levels, displacing apoA-I and reducing HDL CEC, while SAA deletion in mice preserves efflux capacity^37-39^. These studies linking impaired RCT and inflammation, along with the notable enrichment of APR proteins on HDL particles in CVD patients, suggest that impaired RCT due to inflammatory remodeling of the HDL proteome contributes to atherosclerosis progression.

Few studies have investigated the impact of postoperative stress on HDL, RCT, and atherosclerosis. Janssen et al. (2015) first showed in a mouse model that perioperative stress increases plaque volume and vulnerability^40^. This finding was further corroborated by Fuijkschot et al. (2016) and Handke et al. (2021), who observed increased plaque necrosis and size after surgery^41,42^. However, these studies are confounded by the combined effects of blood loss and surgery, as hemodynamic changes by themselves are linked to major adverse cardiovascular events (MACE)^43^. This study isolates the effects of surgery-induced inflammation by utilizing an abdominal laparotomy model that minimizes blood loss and avoids perioperative blood draws. We show that postoperative stress acutely remodels HDL and impairs RCT, leading to rapid lipid accumulation, foam cell apoptosis, and necrotic core expansion in atherosclerotic plaques. Using a novel dual-cell-type RCT model, we demonstrate that macrophage-derived RCT is selectively impaired, while VSMC-derived RCT remains largely intact. Notably, immediate apoA-I supplementation at the time of surgery partially restores RCT and mitigates plaque lipid buildup, suggesting a potential therapeutic strategy to reduce postoperative cardiovascular risk.

## METHODS

### Data availability

The data supporting the findings of this study are included in the article and Supplemental Material. Any additional data will be made available by the corresponding author upon reasonable request.

### Animals

All procedures were approved by the University of Ottawa Animal Care and Use Committee. Male and female *apoE*^*-/-*^ mice (B6.129P2-*Apoe*^*tm1Unc*^/J; strain 002052) were acquired from Jackson Laboratory and kept on a normal laboratory diet (Envigo; 2019) in a temperature and light-controlled environment. The animals were started on Western diet (WD; Envigo; TD.88137, 0.2% cholesterol) at 8 weeks of age to accelerate atherosclerosis development. C57BL/6 mice were obtained from Charles River Laboratories for preliminary studies (C57BL/6N), and from Jackson Laboratories (C57BL/6J) when used as cell donors for reverse cholesterol transport experiments in *apoE*^*-/-*^ animals.

### Abdominal Laparotomy

8-week-old *apoE*^*-/-*^ mice were fed a WD for 8 weeks and subjected to an exploratory abdominal laparotomy. Mice received 1.2 mg/kg buprenorphine SR subcutaneously 1 hour before a 30-minute surgery under 2-3% isoflurane sedation on a warming surface. A 4 cm midline incision was made, intestines were gently manipulated in four locations using a saline-soaked cotton Q-tip (5 strokes/quadrant), before closing with sutures. Fluid support in the form of 200 µL of subcutaneous warm 0.9% saline was provided. Anesthesia-only control mice were similarly sedated and received 200 µL of subcutaneous warm 0.9% saline. Animals were monitered per instituational guidelines following their procedure to confirm appropriate recovery. Animals were randomized into baseline, surgery, or anesthesia control groups based on weights and plasma cholesterol (assessed using the *Infinity*^*TM*^ cholesterol liquid stable reagent, Thermo Scientific).

### Plasma Cytokine Quantification

Plasma SAA from individual animals was assessed using a Mouse SAA ELISA Kit (Invitrogen), as per the manufacturer’s protocol. Other inflammatory cytokines were assessed with the LEGENDplex Mouse Inflammation Panel 13-plex (Biolegend), pooling plasma of 3-5 mice from each condition and running in technical duplicates.

### Lipoprotein profiling

Lipoprotein fractions were obtained through FPLC separation of 200 µL plasma using a Superose 6 Increase column at flow rate 0.75 mL/minute. Following sample injection, 500 µL fractions were collected and assessed for total cholesterol (Pointe Scientific), phospholipids (Fujifilm) and triglycerides (Fujifilm). Lipoprotein fractions were defined as follows: VLDL fractions 3-7, LDL fractions 8-15, HDL fractions 16-20. 200 µL of each HDL fraction was combined to create an HDL pool for HDL proteomics and cholesterol efflux studies.

### Histological Analyses

All histological assessments were conducted in accordance with American Heart Association guidelines^44^. Aortic roots were collected at sacrifice, embedded in OCT, flash frozen and stored at −80°C until sectioning. The aortic sinuses were serially sectioned at 10 µm thickness and spaced 100 µm apart. Sections were stored at −20°C until use. Hematoxylin and eosin (H&E), Oil Red O (ORO), and Mason’s Trichrome (MT) staining were performed to evaluate plaque area, necrotic area (defined as acellular regions), neutral lipid content, and collagen deposition, respectively. Slides were imaged using a Leica Aperio 8 slide scanner with an HC (high capacity) Plan-Apochromat 20×/0.8 objective. Image analysis was performed using the Fiji software^45^.

### Immunophenotyping

50 µL of whole blood was collected by cardiac puncture in EDTA-coated tubes and stained immediately for flow cytometry. Red blood cells were lysed using PharmLyse (BD Biosciences). FC block was performed for 5 minutes (BD Biosciences) in Brilliant Stain Buffer (BD Biosciences). An antibody mastermix was added and incubated for 20 minutes at 4°C, before adding Fixable Viability Stain 700 and incubating for an additional 30 minutes at 4°C. The cells were fixed with Cytofix (BD Biosciences) and analysed on a BD LSRFortessa™. The immune cell populations were defined as follows (**Fig.S1a**): leukocytes CD45^+^; inflammatory monocytes CD11b^+^ Ly6G^-^ Ly6C^hi^; neutrophils CD11b^+^ Ly6C^+^ Ly6G^+^. The analysis was performed using FlowJo (Becton Dickinson). A complete list of antibodies is available in the data supplemental.

### Aortic digest and flow cytometry staining

Mice were perfused with HBSS and the aortic arches and branches (brachiocephalic, left common carotid and left subclavian arteries) were dissected and cleaned of all surrounding tissue. Aortic arches were cut into small pieces, and enzymatically digested with 0.4 units/mL Liberase™ (Roche), 40 units/mL hyaluronidase (Sigma) and 20 units/mL DNAse (Sigma) in HBSS (1.4 µM CaCl_2_, 1 mM EDTA) for 15-20 minutes at 37°C. The cell suspension was passed through 70 µm cell strainer and stained for flow cytometry. FC block was performed for 5 minutes in Brilliant Stain Buffer at RT. An antibody mastermix was added and incubated for 20 minutes at 4°C, before adding Fixable Viability Stain 510 and incubating for an additional 30 minutes at 4°C. The cells were fixed with Cytofix (BD Biosciences). BODIPY staining was performed for 30 minutes at RT and analysed immediately after on a BD LSRFortessa™. The immune cell populations were defined as follows (**Fig.S2**): leukocytes CD45^+^; myeloid cells CD45^+^CD11b^+^. The analysis was performed using FlowJo (Becton Dickinson). A complete list of antibodies is available in the data supplemental.

### Immunohistochemistry

Frozen aortic root sections were air dried for 20 minutes at room temperature (RT) under air flow and fixed in 4% PFA for 10 minutes at RT. Slides were washed thrice for 5 minutes in PBS, then blocked in 10% horse serum + 0.1% Triton X-100 at RT. Primary antibodies were prepared in 1% horse serum + 0.1% Triton X-100, and incubated overnight at 4°C in a moisture chamber. Slides were washed thrice in PBS before adding secondary antibodies (prepared in 1% horse serum + 0.1% Triton X-100), and incubating for 2 hours at RT. Slides were washed thrice in PBS, counterstained with Hoechst 3342, mounted in glass antifade mounting media, and left to dry overnight. Imaging was performed on a Zeiss Axio Observer.Z1 / 7, using a Plan-Apochromat 20x/0.8 M27 objective. Image analysis was performed with the Fiji software, and with the ZEISS V3.11 ZenDesk 2D image analysis toolkit (**Fig.S3a**).

### HDL Proteomics Analysis

#### Protein digestion and peptide reduction

Pooled HDL fractions from 24-hour postoperative mice (n=5) and anesthesia control mice (n=5), isolated as described in *lipoprotein profiling*, were applied to lipid removal agent (LRA, MilliporeSigma™ Supelco™) and washed with 25 mM ammonium bicarbonate to remove non-lipoprotein associated proteins. LRA-bound lipoprotein pellets were incubated overnight at 37°C in 0.05 µg/µL sequencing grade trypsin (Promega) dissolved in 25 mM ammonium bicarbonate. The next morning, samples were vortexed vigorously, centrifugated at 5,000 x *g*, and supernatants (peptide digests) were collected and stored at −80°C. For peptide reduction and alkylation, 20 μg of protein, as determined by protein assay, was transferred to a new tube and the volume was adjusted to 60 μL with 50 mM ammonium bicarbonate (ABC) containing 1% sodium deoxycholate (DOC). Peptide thiols were reduced with 0.2 mM dithiothreitol (DTT) at 37°C for 30 minutes, then alkylated with 0.8 mM iodoacetamide (IAA) at 37°C for 30 minutes in the dark. The sample was acidified, and DOC was precipitated by adding 10 μL of 50% formic acid, then centrifuged at 16,000 x *g* for 5 minutes at RT, and the supernatant was recovered. Peptides were purified using C18 StageTips, dried under vacuum, and resuspended in 21 μL of 2% acetonitrile (ACN) and 0.05% trifluoroacetic acid (TFA). Peptide concentration was estimated using 1 μL for absorbance readings at 205 nm (Nanodrop) and adjusted to 0.2 μg/μL with 2% ACN and 0.05% TFA.

#### LCMS Analysis

The nanoLC-MS/MS system comprised a U3000 NanoRSLC liquid chromatography system (ThermoScientific, Dionex Softron GmbH, Germering, Germany) in line with an Orbitrap Fusion Tribrid–ETD mass spectrometer (ThermoScientific, San Jose, CA, USA), driven by Orbitrap Fusion Tune Application 3.3.2782.34 and equipped with a Nanospray Flex ion source with a FAIMS Pro interface. An equivalent amount of 1 μg of each sample was injected in 5 μL. Peptides were trapped at 20 μL/min in loading solvent (2% ACN / 0.05% TFA) on a 300 mm i.d x 5 mm, C18 PepMap100, 5 mm, 100 Å precolumn cartridge (Thermo Fisher Scientific) for 5 minutes. The pre-column was switched in line with a PepMap100 RSLC, C18 3 mm, 100 Å, 75 μm i.d. x 50 cm length column (Thermo Fisher Scientific), and the peptides were eluted with a linear gradient from 5–40% solvent B (A: 0.1% FA, B: 80% ACN / 0.1% FA) over 90 minutes at 300 nL/min, for a total run time of 120 minutes. Spectra were acquired using Thermo XCalibur software 4.3.73.11. Lock mass internal calibration on the m/z 445.12003 siloxane ion was used. A total cycle time of 3 sec was equally split between three methods with a Compensation Voltage (CV) of either −40V, −50V, or −60V. For each method, full-scan mass spectra (350–1800 m/z) were acquired in the Orbitrap using an AGC target set on Standard, a maximum injection time of 50 ms, and a mass resolution of 120,000. Each MS scan was followed by the acquisition of fragmentation MS/MS spectra of the most intense ions for a total cycle time of 1.5 seconds (top speed mode). The selected ions were isolated using the quadrupole analyzer in a window of 1.6 m/z and fragmented by higher-energy collision-induced dissociation (HCD) with 35% collision energy. The resulting fragments were detected in the ion trap with a normalized AGC target of 33% and a maximum injection time of 50 ms. Dynamic exclusion of previously fragmented peptides was set for a period of 30 sec with a tolerance of 10 ppm. This protocol ensured efficient HDL proteome characterization with high specificity and sensitivity using nanoLC-MS/MS.

#### Spectral annotation and label-free protein quantification

Mass spectrometry data processing and relative quantification was performed using Proteome Discoverer (Thermo Scientific, version 3.1.0.638). Raw files were searched against the UniProt *Mus musculus* protein database (version 2024-01-24) using Sequest HT. Search parameters included “trypsin” as the digestion enzyme with a maximum of 2 missed cleavage sites, carbamidomethylation of cysteine was set as a fixed modification, methionine oxidation and N-terminus protein acetylation were set as variable modifications. Mass tolerances were 10 ppm and 0.5 Da for MS and MS/MS, respectively. Peptide and protein identifications were filtered at 1% False Discovery Rate (FDR). Label-free quantification was performed based on precursor ion intensities. Only proteins with at least 2 unique peptides and detectable intensity values in 75% of samples in at least one group were considered as quantifiable proteins. Volcano plots were generated by plotting log_2_ fold change vs −log_10_(p-value). Heatmap of differentially expressed proteins identified in both groups was generated using Heatmapper^46^, and includes proteins meeting the thresholds of an FDR-adjusted q-value of < 0.05 and a |Z score| > 1.96.

#### Go Biological Pathway

GO Biological Pathways were extracted using the ShinyGo 0.80 tool^47^. HDL-associated proteins meeting a statistical significance of P < 0.05 from surgery (upregulated after surgery) or anesthesia-only (downregulated after surgery) groups were fed to the program with the following specifications: FDR cutoff: 0.05; pathway min size: 15 max size: 1000. Statistical significance of included proteins was chosen to capture a broader range of differentially abundant proteins, and regulated pathways of interest were subsequently evaluated with functional assays (e.g. *in vitro* cholesterol efflux).

### Cell culture

Murine bone marrow–derived macrophages (BMDMs) were generated by harvesting bone marrow from the long bones of C57Bl/6N mice and culturing for 7 days in DMEM media supplemented with 20% L929 conditioned media (made in-house), 10% fetal bovine serum (FBS), and 1% penicillin-streptomycin. Cells were maintained at 37°C and 5% CO_2_. Primary murine VSMCs were generated as previously described^13^, and used at passage 3-5.

### Reverse Cholesterol Transport

For in vivo RCT assays, radiolabeled-cholesterol-loaded cells were injected subcutaneously into mice post-surgery, with plasma, liver, bile, and feces assessed for cholesterol movement, as previously^48^. 8-week-old male *apoE*^-/-^ mice fed a WD for 8 weeks were randomized into surgery or anesthesia-only groups. Aggregated LDL (agLDL, 50 μg/mL) was prepared using endotoxin-free LDL from human plasma and incubated with [^3^H]-cholesterol (5 μCi/mL) for 1 hour at 37°C. Bone marrow-derived macrophages (BMDMs) from age-matched mice were incubated with radiolabeled agLDL for 30 hours in 10% FBS-containing DMEM media, washed with HBSS, and equilibrated overnight in 2 mg/mL fatty acid-free BSA. Cells were then washed in ice-cold HBSS, incubated with 5 mM EDTA for 20 minutes at 4°C, spun at 400 x *g* for 5 minutes, resuspended in ice-cold DMEM, and injected subcutaneously into mice before waking from anesthesia. After surgery, plasma (24 hours and 48 hours), liver (48 hours), bile (48 hours), and feces (48 hours) were collected and analyzed for radioactivity on a Perkin Elmer 4810TR Tricarb liquid scintillation counter.

RCT from VSMCs was similarly performed; VSMCs were loaded for 48 hours with methyl-β-cyclodextrin (mβCD)-cholesterol (35 μg/mL) and [^3^H]-cholesterol (5 μCi/mL) in DMEM media containing 10% FBS, equilibrated overnight in 2 mg/mL fatty acid-free BSA, lifted in trypsin (0.05%) - EDTA (0.002%), spun at 400 x *g* for 5 minutes, resuspended in ice-cold DMEM, and injected subcutaneously into mice before waking from anesthesia. RCT was assessed as above. For dual-label RCT, BMDMs and VSMCs were incubated with mβCD-cholesterol (35 μg/mL), with [^3^H]-cholesterol (VSMCs; 5 μCi/mL) or [^14^C]-cholesterol (BMDMs; 1 μCi/mL) for 48 hours at 37°C, before incubating overnight in 2 mg/mL fatty acid-free BSA. BMDMs and VSMCs were lifted and prepared into separate injections in ice cold DMEM. The cells were injected subcutaneously sequentially during anesthesia recovery, with BMDMs in the scruff of the neck and VSMCs 3-cm lower. CPMs and DPMs were measured from tissues as above on a Perkin Elmer 4910TR Tricarb liquid scintillation counter with dual radiotracer detection capacity.

### Cholesterol efflux

Cholesterol efflux experiments were performed as previously described^49^. Briefly, BMDMs or VSMCs were incubated for 48 hours with ^3^H-cholesterol (5 µCi/mL), then incubated overnight in DMEM containing 2 mg/mL of fatty acid-free BSA. The next morning, the media was replaced with 2 mg/mL of fatty acid-free BSA DMEM with or without 1% mouse plasma or 5% isolated HDL (FPLC-isolated as above; 5% equivalent to an average of 50 µg/mL HDL across all conditions). Plasma and HDL were filter sterilized using 0.22 µm Costar® Spin-X® centrifuge tube filters beforehand. Cholesterol acceptors were incubated for 6 or 24 hours. BSA media containing recombinant ApoA-I (produced in house) (50 µg/mL) was used as a positive control. After incubation, the supernatants were collected and spun at 1000 x *g* for 5 minutes to remove cell debris. The cells were washed thrice with PBS and lysed in 0.5 M NaOH. Supernatants and cell lysates were read on a Perkin Elmer 4810TR Tricarb liquid scintillation counter. Cholesterol efflux is expressed as a percentage of ^3^H-cholesterol in the supernatant/(^3^H-cholesterol in the supernatant + ^3^H-cholesterol in cells) x 100%. Plasma-specific and HDL-specific efflux were calculated by subtracting % cholesterol efflux in 2 mg/mL BSA media-only control wells.

### Cholesterol efflux to postoperative human plasma

Blood from male and female patients undergoing cancer surgery at The Ottawa Hospital was collected from patients on the morning of their procedure as part of an REB approved research protocol (OHSN REB-2011884-011H). The samples were collected in sodium heparin tubes and spun at 1000 x *g* for 10 minutes. The upper plasma layer was collected and centrifuged again at 10 000 x *g* for 15 minutes. Supernatants were collected and stored at −70°C. To assess cholesterol efflux, THP-1 cells were incubated for 48 hours with ^14^C-cholesterol (1 µCi/mL) in 10% FBS RPMI media, then equilibrated overnight in RPMI containing 2 mg/mL of fatty acid-free BSA. The next morning, the media was replaced with 2 mg/mL fatty-acid free RPMI media containing 2% human plasma, incubated for 6 hours, and collected / analyzed as described in *Cholesterol Efflux* methods section. Plasma samples were filter sterilized beforehand using 0.22 µm Costar® Spin-X® centrifuge tube filters.

### Western Blotting

4X Laemmli Sample Buffer (Bio-Rad) containing β-mercaptoethanol was added to 3 µL plasma and boiled at 95°C for 5 minutes. Samples were run on 8-16% Criterion TGX Stain Free Pre-cast Gels. Proteins were transferred onto 0.45 µm PVDF membranes using the Trans-Blot Turbo Transfer System (Bio-Rad). Immunoblotting was performed with primary antibodies at RT for 2 hours, followed by horseradish peroxidase conjugated secondary antibodies for 1 hour at RT. Proteins were developed using Clarity (Bio-Rad) Substrates and imaged on a ChemiDoc MP system (Bio-Rad).

### ApoA-I intervention

Human recombinant ApoA-I was synthesized and purified as previously described^50^. For studies intervening with ApoA-I, 40 mg/kg ApoA-I diluted in saline was injected intraperitoneally in a 200 µL final volume at 0 hours (upon anesthesia recovery) and 24 hours postoperative.

### Statistical Analyses

Data are represented as mean ± SEM. Statistical tests were performed on GraphPad Prism V10.3.1 software (GraphPad Software, Inc) using 2-way ANOVA with Holm-Sidak multiple comparison test, or unpaired 2-tailed t test. Outliers were identified and removed using the ROUT (Robust Regression and Outlier Removal) method (Q=1%).

## RESULTS

### Rapid Necrotic Core Expansion in Atherosclerotic Plaques Following Surgery

Observational studies report a spike in cardiovascular events after major surgery, with increased age, male sex, and atherosclerosis as independent risk factors. However, the mechanisms of postoperative atherosclerotic plaque destabilization remain unclear. Systemic inflammation from surgery rapidly accelerates atherosclerosis in mice^40-42^, but these findings are confounded by the combined effects of blood loss and surgery, as hemodynamic changes alone are linked to MACE^43^. To isolate the impact of surgery-induced inflammation, we developed an abdominal laparotomy model with minimal blood loss and no perioperative blood draws, preserving blood volume and excluding blood loss as a confounder. In C57BL/6 male mice subjected to abdominal laparotomy, our surgical model induced an APR, with serum SAA increasing 1000-fold, IL-1β 200-fold, and IL-6 300-fold post-surgery, progressively returning to baseline at 48h (**Fig.1a-d**).

To determine the impact of surgery-induced inflammation on atherosclerosis using this surgery model, 8-week-old male and female *apoE*^*-/-*^ mice were placed on WD for 8 weeks and randomized into baseline, 24h, 72h, and 15-day post-surgery groups (**Fig.1a, e, f**). Lipoprotein profiles at 24h post-surgery revealed a decrease in VLDL cholesterol and phospholipids, likely related to reduced oral intake in the immediate postoperative period (**Fig.1g-i**). No significant differences were observed in the lipid composition of LDL or HDL, although small changes in HDL lipid composition were likely masked by low baseline levels in *apoE*^*-/-*^ mice (**Fig.1g-i**). Atherosclerotic plaques were quantified for size at the aortic root (**Fig.1k**). While surgery alone did not alter lesion area in the aortic sinus (**Fig.1l**), the necrotic core area was significantly expanded 15 days after surgery (**Fig.1m**). Collagen deposition did not differ between the surgery and anesthesia-only control groups (**Fig.1n**). Intriguingly, a decrease in Oil Red O (ORO) staining, indicative of neutral lipids, was observed in surgery mice 15 days postoperatively (**Fig. 1o**).

**Fig 1:**
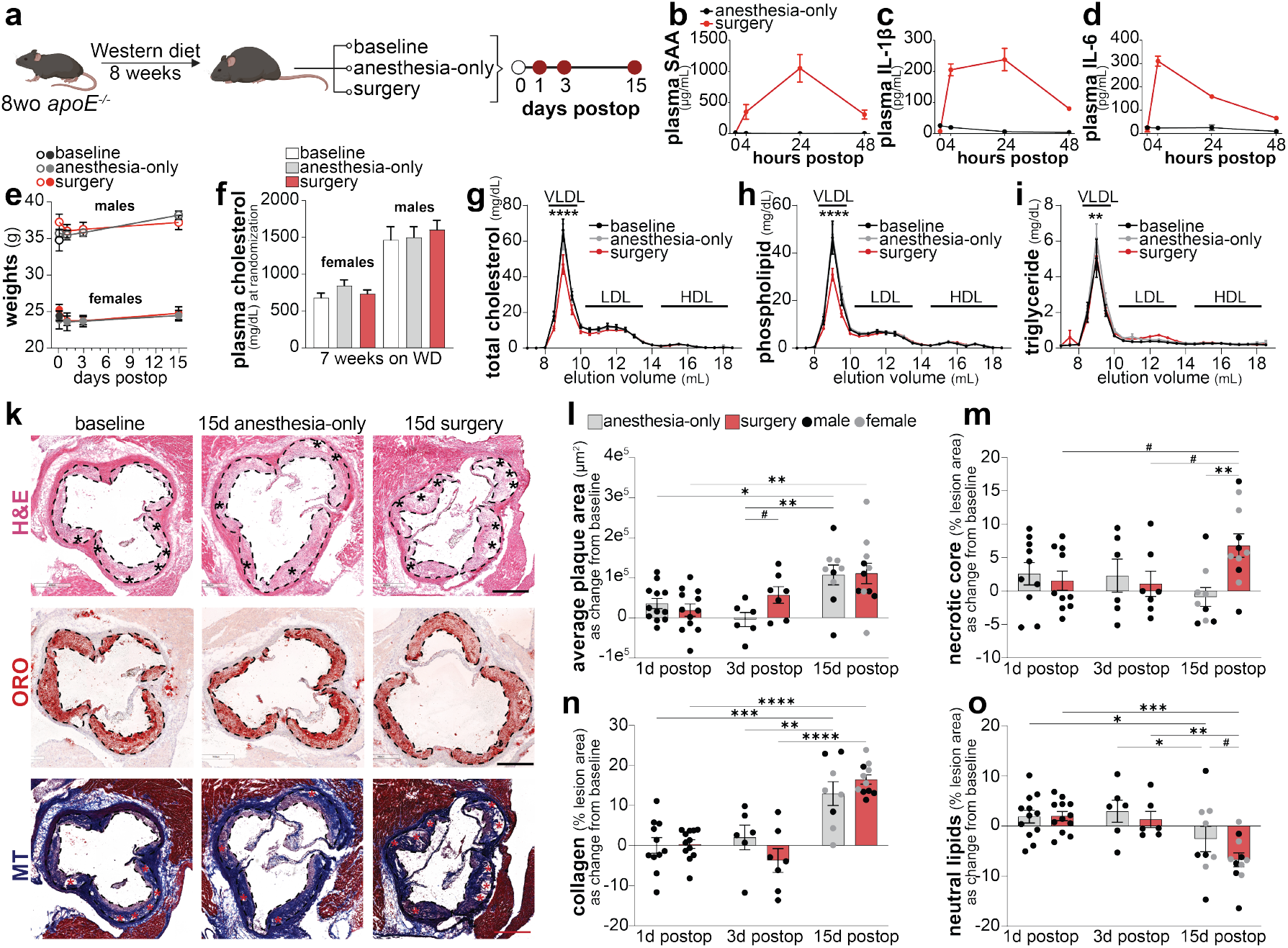
Surgery drives necrotic core expansion. 8-week-old *apoE*^*-/-*^ mice (males and females) were fed a Western diet for 8 weeks and randomised into baseline (n=12), anesthesia control (n=6-12/timepoint), or surgery groups (n=6-11/timepoint) (**a**). (**b-d**) 12-week-old male C57BL/6 mice underwent abdominal laparotomy (n=5) or anesthesia (n=3). Levels of plasma SAA (**a**), IL-1β (**b**), and IL-6 (**c**) at baseline, 4, 24, & 48h were quantified. Mouse weights (**e**) and plasma cholesterol at randomization (**f**). Total cholesterol (**g**), phospholipids (**h**), and triglycerides (**i**) of HPLC-fractionated plasma from 24h postoperative and controls (n=6/group). H&E-, Oil Red O (ORO)-, and Masson’s Trichrome (MT)-stained atherosclerotic plaques in the aortic root (necrotic regions labelled with stars) (**k**). Average plaque area (**l**), necrotic core (**m**), collagen (**n**), and neutral lipid (**o**), as absolute change from the baseline plaques. Data presented as mean ± SEM. # P ≤ 0.1, * P ≤ 0.05, ** P ≤ 0.01, *** P ≤ 0.001, **** P ≤ 0.0001.

### Acute Lipid Accumulation in Arterial CD45^+^ and CD45^-^ Cells After Surgery

Plaque destabilization due to necrotic core expansion can result from various mechanisms, including immune cell infiltration, inflammation, and lipid deposition. As expected in the postoperative period, we found increased circulating neutrophils and Ly6C^hi^ inflammatory monocytes (**Fig.2a**), two pro-atherogenic cell populations. To determine whether circulating inflammatory immune cells infiltrated plaques, increasing inflammation and cellularity, and to evaluate lipid loading and foam cell formation in situ, atherosclerotic plaques were assessed in two disease-prone regions of the aorta: the arch and the aortic sinus. In the arch, a striking increase in neutral lipids (per BODIPY staining) was found in the aortic myeloid cells of 24h post-surgery mice as compared to controls (**Fig.2b**).

**Fig 2:**
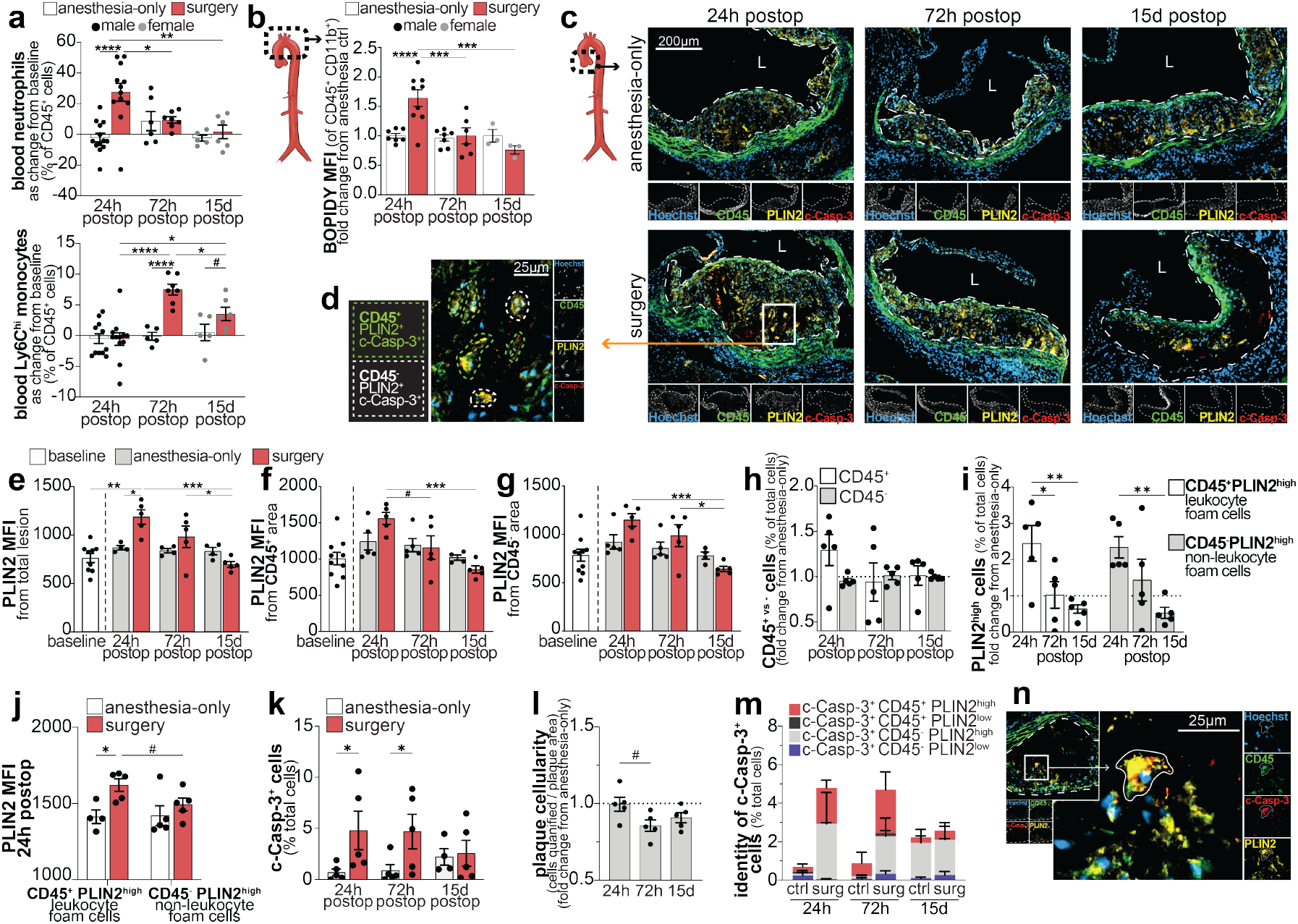
Surgery increases lipid buildup in foam cells of atherosclerotic plaques. Circulating neutrophils (CD45^+^CD11b^+^Ly6G^+^) and inflammatory monocytes (CD45^+^CD11b^+^Ly6G^-^Ly6C^hi^) in anesthesia-only and postoperative mice by flow cytometry (n=5-12/group/timepoint) (**a**). BODIPY MFI in myeloid cells (CD45^+^CD11b^+^) of aortic digests by flow cytometry, as fold change from anesthesia control (**b**). Representative images of aortic root IF (Hoechst, CD45, PLIN2, c-Casp-3) (n=5/group) (**c**), with representation of CD45^+^ (green outline) vs CD45^-^ (white outline) foamy apoptotic cells (**d**). PLIN2 MFI in total (**e**), CD45^+^ (**f**), and CD45^-^ (**g**) plaque areas. Per cell analysis of IF images (**h-m**): percentage of total plaque cells of CD45^+^ and CD45^-^ cells, as fold change from anesthesia-only control (**h**). CD45^+^PLIN2^high^ and CD45^-^PLIN2^high^ cells, as percentage of total plaque cells (**i**). PLIN2 MFI of CD45^+^PLIN2^high^ and CD45^-^PLIN2^high^ cells 24h after surgery (**j**). c-Casp-3^+^ plaque cells (percentage total cells) (**k**). Plaque cellularity (number of cells quantified / plaque area) after surgery. as change from anesthesia control (**l**). CD45 and PLIN2 expression of c-Casp-3^+^ cells; cell populations shown as percentage of total plaque cells (**m**). Representative image of a c-Casp-3^+^CD45^+^PLIN2^high^ cell in the plaque (**n**). Data presented as mean ± SEM. # P ≤ 0.1, * P ≤ 0.05, ** P ≤ 0.01, *** P ≤ 0.001, **** P ≤ 0.0001.

Perilipin 2 (PLIN2), a protein found on the surface of lipid droplets that increases in expression as foam cells accumulate lipids^51^, also increased in plaques of postoperative mice (**Fig.2c-e**). Notably, PLIN2 expression was elevated in both the CD45^+^ (**Fig.2f**) and CD45^-^ (**Fig.2g**) plaque areas of the aortic sinus 24h after surgery. Using the ZEISS Zendesk 2D image analysis toolkit, we quantified the per-cell expression of CD45, PLIN2 and cleaved (active) caspase-3 (c-Casp-3), an apoptosis marker (**Fig.2h-m, Fig.S3**). Foam cell sub-population analysis revealed a small influx of CD45^+^ cells 24h after surgery (**Fig.2h**), and increased PLIN2 expression in both CD45^+^ (leukocyte) and CD45^-^ (non-leukocyte) foam cells 24h after surgery (**Fig.2i**), with leukocyte foam cells showing higher PLIN2 expression (**Fig.2j**). Elevated numbers of c-Casp-3^+^ cells were found in plaques 24h and 72h after surgery (**Fig.2k**), correlating with decreased plaque cellularity at 72h and 15 days post-surgery (**Fig.2l**). Apoptotic cells were of both leukocytic and non-leukocytic origin and almost exclusively PLIN2^high^ (**Fig.2m,n**). Taken together, our findings reveal the remarkably rapid dynamics of lipid droplets in plaque foam cells during the APR and suggest that the exacerbation of lipid buildup in macrophage and VSMC foam cells after surgery drives necrotic core expansion.

### Postoperative Inflammatory Remodeling of HDL

Next, we used label-free proteomics to analyze HDL proteomes of 24h postoperative mice and anesthesia controls. Of the 392 proteins identified on HDL, 140 were differentially abundant (P < 0.05; 100 more abundant in surgery, 40 more abundant in control). Principal component analysis distinguished the groups (**Fig.3a**), with a notable increase in SAA1 and SAA2 after surgery and reduced apoA-I (**Fig.3b**). Postoperative HDL was highly enriched with proteins related to the APR and acute inflammation, as indicated by significant Z-scores (|Z| > 1.96) and GO Biological Process terms (P < 0.05) (**Fig.3c,d**). Conversely, pathways related to RCT such as cholesterol efflux and lipid transport pathways were the most downregulated in postoperative HDL (**Fig.3d**). Notably, cholesterol efflux from BMDMs and VSMCs to whole plasma or HPLC-isolated HDL was significantly reduced when using samples from 24-hour postoperative mice compared to those from anesthesia-only controls (**Fig.3e,f**). This phenomenon was also observed in samples obtained from patients undergoing abdominal surgery for cancer, where we observed a striking reduction in cholesterol efflux to postoperative human plasma (**Fig.3g**). These findings underscore a critical disruption in HDL efflux capacity following non-cardiac surgery, highlighting a potential mechanistic link between postoperative inflammation and impaired cholesterol transport, which could have profound implications for cardiovascular health in the postoperative setting.

**Fig 3:**
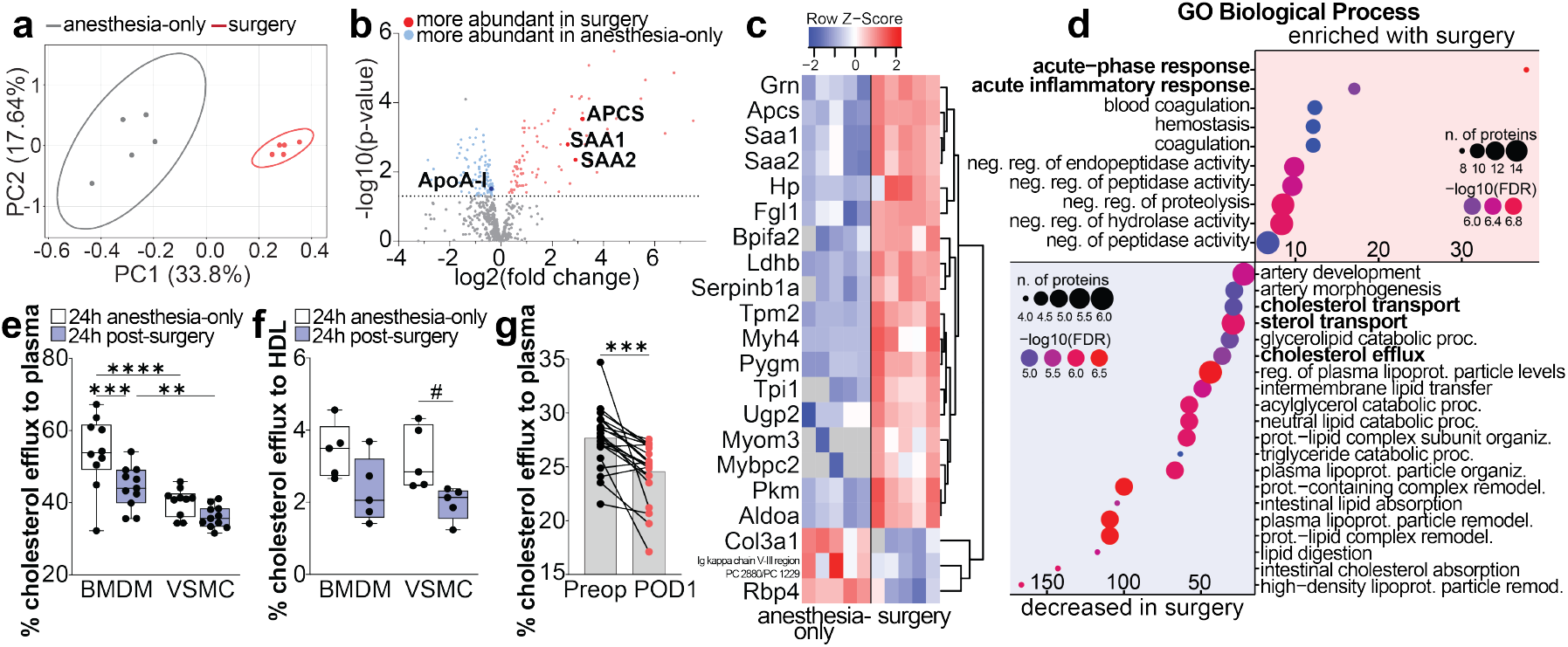
Surgery drives inflammatory remodeling of HDL, reducing cholesterol efflux capacity. HDL was isolated by HPLC from plasma of 24h postoperative and anesthesia control mice (n=5/group). The HDL proteome was analyzed by label-free LC-MS/MS. Principal component analysis of identified proteins (**a**). Volcano plot showing proteins with significantly different abundance (P < 0.05) (**b**). Heatmap of most differently abundant proteins (Limma q-value < 0.05, |Z score| > 1.96) (**c**). GO Biological Processes of proteins significantly enriched on postoperative HDL (10 pathways presented) and enriched on anesthesia control (decreased in surgery; 20 pathways presented) (**d**). 24h cholesterol efflux of BMDMs and VSMCs to 1% plasma from 24h postoperative or anesthesia control mice (n=10-11/group) (**e**). 6h cholesterol efflux of BMDMs and VSMCs to 5% isolated HDL (∼50µg/mL) from 24h postoperative or anesthesia control mice (n=5/group) (**f**). ^14^C-cholesterol efflux to preoperative (preop) or postoperative day 1 (POD1) plasma (2%) of patients undergoing general surgery (n=21; lines connect results from one patient) (**g**). Data as min to max boxplot, # P ≤ 0.1, ** P ≤ 0.01, *** P ≤ 0.001, **** P ≤ 0.0001 (**e,f**). Unpaired two-tailed T-test, ***: P < 0.0001 (**g**).

### Postoperative Impairment of Reverse Cholesterol Transport

To quantify RCT *in vivo*, 8-week-old male *apoE*^*-/-*^ mice fed a WD for 8 weeks were randomized into surgery or anesthesia-only groups. ^3^H-cholesterol-loaded macrophages were injected subcutaneously before waking from anesthesia. Afterward, cholesterol movement was assessed in the plasma, liver, bile, and feces, as previously^48^. We found a marked reduction in ^3^H-cholesterol movement from BMDMs into plasma, liver, and feces after surgery (**Fig.4a**). Likewise, in wild-type C57BL/6 mice, macrophage RCT was significantly reduced after surgery (**Fig.S4**).

**Fig 4:**
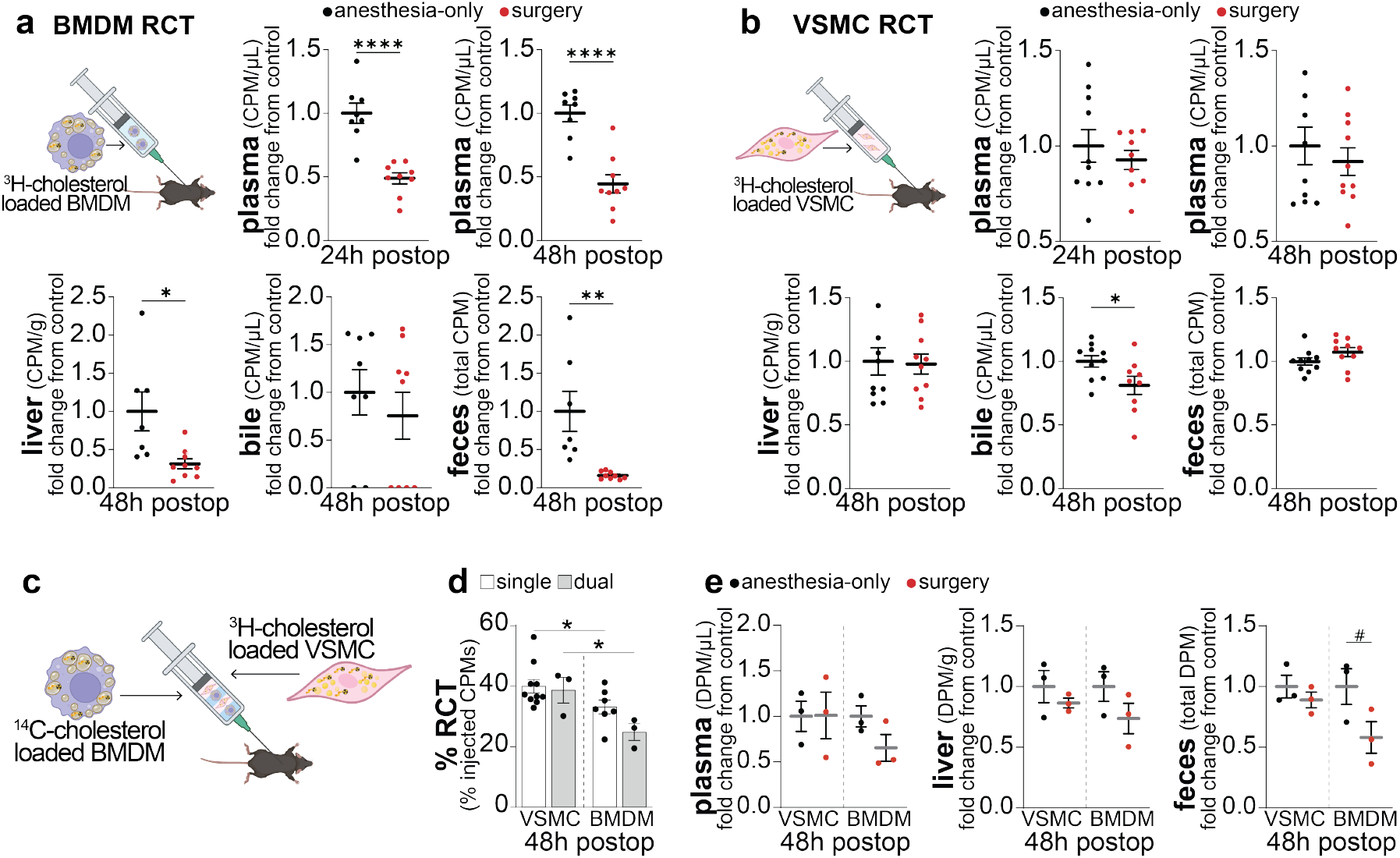
Surgery impairs macrophage RCT. Male *apoE*^*-/-*^ mice (n=8-9/males group) fed a Western diet for 8 weeks were randomized to anesthesia control or surgery group. BMDMs from sex-& aged-matched C57BL/6 donors were loaded with ^3^H-cholesterol-labeled agLDL for 30h and injected during recovery from anesthesia. Radioactivity was measured in plasma at 24h after surgery, and in plasma, in liver & feces at 48h postoperative (**a**). Similarly, VSMCs loaded with ^3^H-cholesterol and mβCD-cholesterol for 48h were injected in mice to assess postoperative VSMC RCT (n=10 males/group) (**b**). Experimental design for proof-of-concept for *in vivo* dual RCT study, cells were loading with mβCD-cholesterol and radiolabeled cholesterol (^14^C for BMDMs, ^3^H for VSMCs) for 48h before injecting postoperatively (**c**). Total RCT was assessed in anesthesia controls from single-label, single-cell studies (CPMs in quantified tissues/injected CPMs) and a dual-label pilot study (16-wk-old ½ male, ½ female *apoE*^*-/-*^, n=3) (**d**). VSMC and BMDM RCT was assess simultaneously in anesthesia control and postoperative mice (16-wk-old ½ male, ½ female *apoE*^*-/-*^, n=3/group). ^3^H DPMs (VSMCs) and ^14^C DPMs (BMDMs) of 48h plasma, liver and feces are shown. Data as mean ± SEM, # P ≤ 0.1, * P ≤ 0.05, ** P ≤ 0.01, **** P ≤ 0.0001.

While macrophage RCT has been well described in mice, VSMC RCT remains largely understudied^52^. Interestingly, VSMC RCT appeared to be minimally affected by surgery, with only slight trends toward reduced cholesterol transport to plasma and bile (**Fig.4b**). To directly compare the RCT from both foam cell subtypes *in vivo*, we developed a novel dual-label, dual-cell-type RCT model. Macrophages and VSMCs were incubated with mβ-CD-cholesterol and either ^14^C-cholesterol (for BMDMs) or ^3^H-cholesterol (for VSMCs) for 48h. This loading process effectively transforms VSMCs into macrophage-like foam cells^53^, allowing for equivalent lipid loading in both cell types^13^. Cells were then injected into recipient mice, and total macrophage (^14^C CPMs) and VSMC (^3^H CPMs) RCT were quantified by summing ^3^H and ^14^C CPMs across all tissues, expressing them as a percentage of total injected CPMs. Results from the dual-label, dual-cell RCT model showed comparable RCT to the single-label model, confirming significant VSMC RCT *in vivo* (**Fig.4d**). Once again, surgery induced significant impairment in BMDM RCT, while VSMC RCT remained largely unaffected (**Fig.4e**).

### Recombinant ApoA-I Mitigates Postoperative RCT Dysfunction

After surgery, we observed marked reductions in plasma apoA-I levels concomitant with increased SAA levels increased in *apoE*^*-/-*^ mice (**Fig.5a-d**). To test if restoring apoA-I could rescue impaired RCT post-surgery, we conducted an interventional study where mice were randomized into three groups: anesthesia-only control + IP saline, surgery control + IP saline, or surgery + IP rApoA-I (40 mg/kg in saline). IP injections were given upon waking from anesthesia and at 24h post-surgery. rApoA-I injection led to increased plasma apoA-I at 24h and 48h post-surgery, far exceeding anesthesia-only control levels (**Fig.5e**). Raising apoA-I post-surgery modestly increased radioactive counts in the bile, liver, and feces (**Fig.5f-j**). Moreover, we observed reduced lipid accumulation in myeloid cells of the aortic arch with apoA-I (**Fig.5k**). Collectively, our results show that HDL inflammatory remodeling impairs RCT, while raising apoA-I levels may help mitigate necrotic core expansion due to lipotoxicity following surgery.

**Fig 5:**
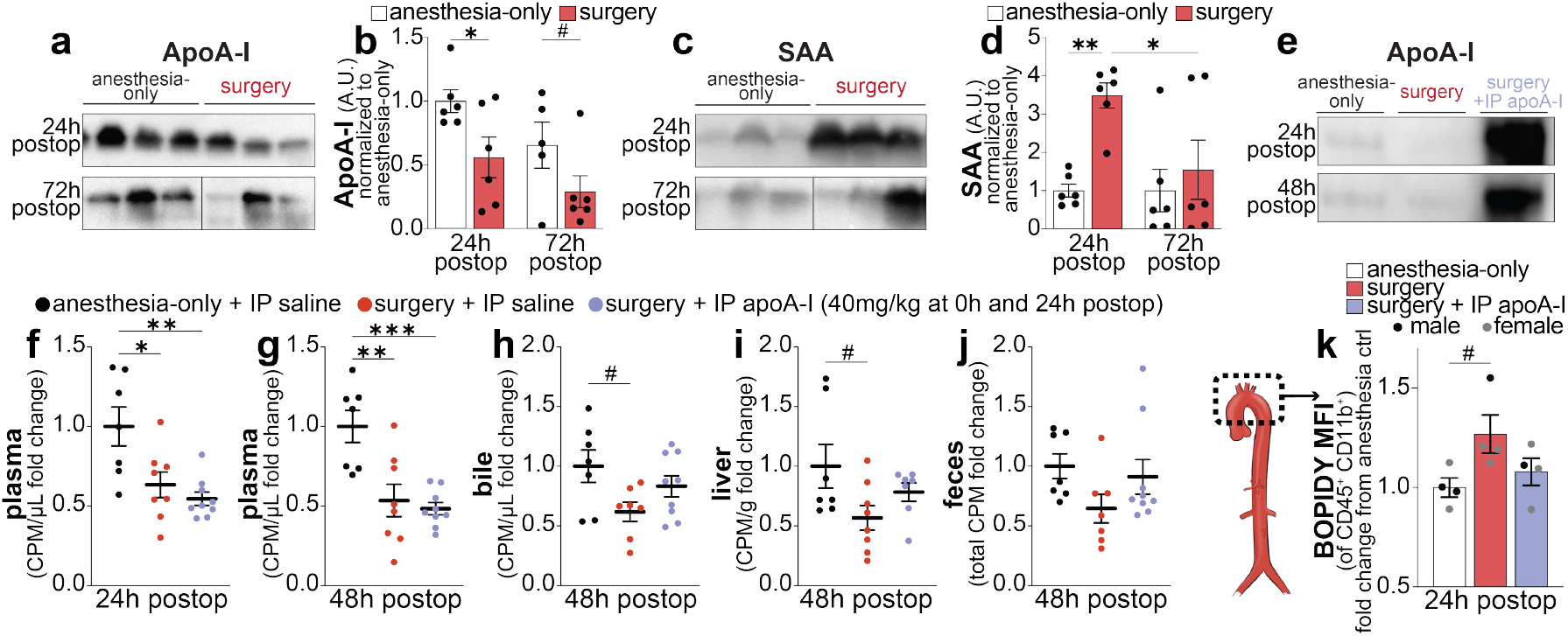
Restoring postoperative apoA-I increases RCT. Western blot of plasma apoA-I (**a,b**) and SAA1+2 (**c,d**) (n=6/group; 3 shown), quantified and normalized to anesthesia control (**b, d**). Plasma apoA-I at 24h & 48h postop in mice injected IP with rApoA-I (40mg/kg) or saline at 0 and 24h postop (**e**). BMDM RCT was assessed with IP rApoA-I injection at 0h and 24h postop, or equivalent volume of saline for control animals, with radioactivity measured as in **Fig. 5a** (n=7-9 males/group) (**f-j**). Aortas from 24h postop mice ± IP rApoA-I were digested, stained for CD45, CD11b, and BODIPY, with BODIPY MFI of CD45^+^CD11b^+^ cells normalized to anesthesia-only controls (n=4/group, ½ males, ½ females) (**k**). Data as mean ± SEM, # P ≤ 0.1, * P ≤ 0.05, ** P ≤ 0.01, *** P ≤ 0.001.

## DISCUSSION

The clinical outcome of atherosclerosis hinges on the balance between pro-inflammatory and inflammation-resolving mechanisms^14^. While lipid accumulation and inflammation drive plaque progression and MACE, inflammation-resolving mechanisms help stabilize plaques, as shown by reduced MACE in trials involving IL-1β antagonism and colchicine^7,54-56^. HDL contributes to cardiovascular health through its anti-inflammatory effects, including cholesterol removal from foam cells and inhibiting pro-inflammatory pathways and cytokines such as TNFα and IL-6^35,57^. While research traditionally focuses on macrophage efflux mechanisms in atherosclerosis, recent studies underscore the importance of studying VSMC-derived foam cells, which constitute the majority of foam cells in plaques^9,58,59^. Therefore, understanding cholesterol efflux in VSMC-derived foam cells is as crucial as in macrophage foam cells.

In this study, we demonstrate that postoperative stress exacerbates lipid accumulation in both macrophage and VSMC foam cells. We find that *in vitro* cholesterol efflux to isolated HDL was similarly reduced in both VSMCs and macrophages, but the reduction in macrophage cholesterol efflux to postoperative plasma was more pronounced than in VSMCs. This reduction in macrophage cholesterol efflux to plasma translated to significant postoperative impairment in RCT, while only modest differences were noted in VSMC RCT. It is possible that the low baseline HDL concentration in *apoE*^*-/-*^ mice masked some differences in postoperative VSMC RCT, while a significant reduction in apoA-I production at the liver may underly the pronounced RCT impairment observed in BMDMs. Indeed, VSMCs primarily rely on ABCG1-mediated cholesterol efflux and preferentially efflux to mature HDL, whereas macrophages promote both ABCA1-and ABCG1-mediated efflux^13^.

Acute lipid accumulation in atherosclerotic plaques was observed in foam cells of both leukocyte and non-leukocyte origin. Leukocyte foam cells accumulated more lipids, which paralleled the impairment in RCT observed in these cell types. Moreover, macrophage-VSMC crosstalk likely occurs within the plaque -cholesterol trafficking to the ER in macrophages treated with atherogenic LDL or with ABCA1/ABCG1 deficiency leads to activation of the NLRP3 inflammasome and IL-1β production^60,61^. Macrophage IL-1β signaling promotes VSMC trans-differentiation, inflammation and apoptosis^62^. This may also contribute to neutrophil recruitment to atherosclerotic plaques and subsequent NETosis^61^. Given the substantial increase in circulating neutrophils 24h after surgery, it is likely that an influx of neutrophils occurs in the early postoperative period, fueling plaque destabilization. Thus, acute cholesterol accumulation in myeloid cells may promote plaque destabilization through cellular crosstalk.

While the capacity of VSMCs for RCT may not be as profoundly impaired as that of macrophages in *apoE*^*-/-*^ mice, VSMCs remain a therapeutic target due to the increased number of foam cells in plaques. Raising postoperative apoA-I shows promise in enhancing postoperative macrophage RCT and reducing acute lipid accumulation, although HDL-based interventions may be preferable to ensure adequate RCT support for both foam cell subtypes.

Uniquely, we observe that metabolic changes induced by postoperative inflammation cause striking and rapid changes in foam cell dynamics within atherosclerotic plaques. Due to the chronic nature of atherosclerosis, studies typically focus on long-term changes, but our findings show that acute systemic disturbances can induce cellular changes in plaque cells as early as 24h postoperative. This highlights the fact that the plaque microenvironment is not shielded from large systemic changes. Furthermore, this observation offers insights into the limited efficacy of clinical trials such as the CSL112 post-MI study, where interventions were initiated up to 5 days after the physiological insult^63,64^. In our model, lipid accumulation and foam cell apoptosis were underway at 24h and 72h postoperative, with significant changes in cellularity at 72h, underscoring a critical therapeutic window that may have been missed.

Previous studies on postoperative atherosclerosis have found success in interventions that temper inflammation (e.g, statins, IL-6 antagonism, preoperative regulatory T cell expansion) and lipid metabolism (e.g. statins)^40,42^. However, our comparison to these studies is limited by the site of atherosclerotic plaque assessment. In those studies, plaque development in the brachiocephalic artery was minimal before surgery, and atherosclerotic plaque formation was primarily the result of the surgical procedure^40,42^. Our results align with those of a study on postoperative atherosclerosis in the aortic root, where an increase in necrotic core was found 15 days after orthopedic surgery^41^. Additionally, in this study, caspase-3 area was not different between the surgery and control groups at 5 and 15 days postoperative, supporting our findings that apoptosis is not altered at 15 days postoperative and that decreased apoptotic signaling likely occurs near the 5-day postoperative timepoint^41^.

Importantly, we replicate our findings of reduced cholesterol efflux to postoperative *apoE*^*-/-*^ plasma in non-cardiac surgery patient samples, further substantiating the translatability of our observations. Recent untargeted metabolomics of patients with and without myocardial injury after non-cardiac surgery (MINS) identified cholesterol metabolism as the KEGG pathway, reinforcing the critical role of lipid metabolism in postoperative cardiovascular complications^65^. Although advances in perioperative medicine have led to improved patient care and safer procedures, cardiovascular complications remain a significant concern, particularly in light of rising cardiovascular disease morbidity. A recent prospective study found that 1 in 6 patients undergoing major general surgery experienced MINS, with MINS associated with a 5-fold increase in 30-day mortality^66^. By understanding the acute changes in foam cell dynamics and cholesterol efflux during the postoperative period, we can better target therapeutic interventions aimed at preserving plaque stability and mitigating the risk of major adverse cardiovascular events (MACE) following surgery. With further investigation into enhancing reverse cholesterol transport, particularly through HDL-based therapies, we may be able to significantly reduce the incidence of postoperative cardiovascular complications, ultimately improving patient outcomes and reducing the burden of MACE in the surgical population.

## ACKNOWLEDGMENTS

The authors thank the University of Ottawa Heart Institute Animal Care and Veterinary Services for technical assistance for mouse atherosclerosis studies, and Megan Fortier for assisting with the surgical procedures. They thank Dr. Vera A. Tang and the University of Ottawa Flow Cytometry & Virometry (FCV) Core Facility for their assistance with flow cytometry experiments. They thank Dr. Rebecca Auer’s lab member for the collection and processing of general surgery patient samples and coordination of sample sharing for cholesterol efflux studies. They thank Dr. Warren Lee and Karen Fung (St. Michael’s Hospital) for providing the recombinant human APOA1 plasmid and for sharing their recombinant APOA1 purification protocol. They thank the University of Ottawa Heart Institute High Resolution Cell Imaging Core Facility for their help with aortic sinus histology.

---

**Supplemental Fig 1:**
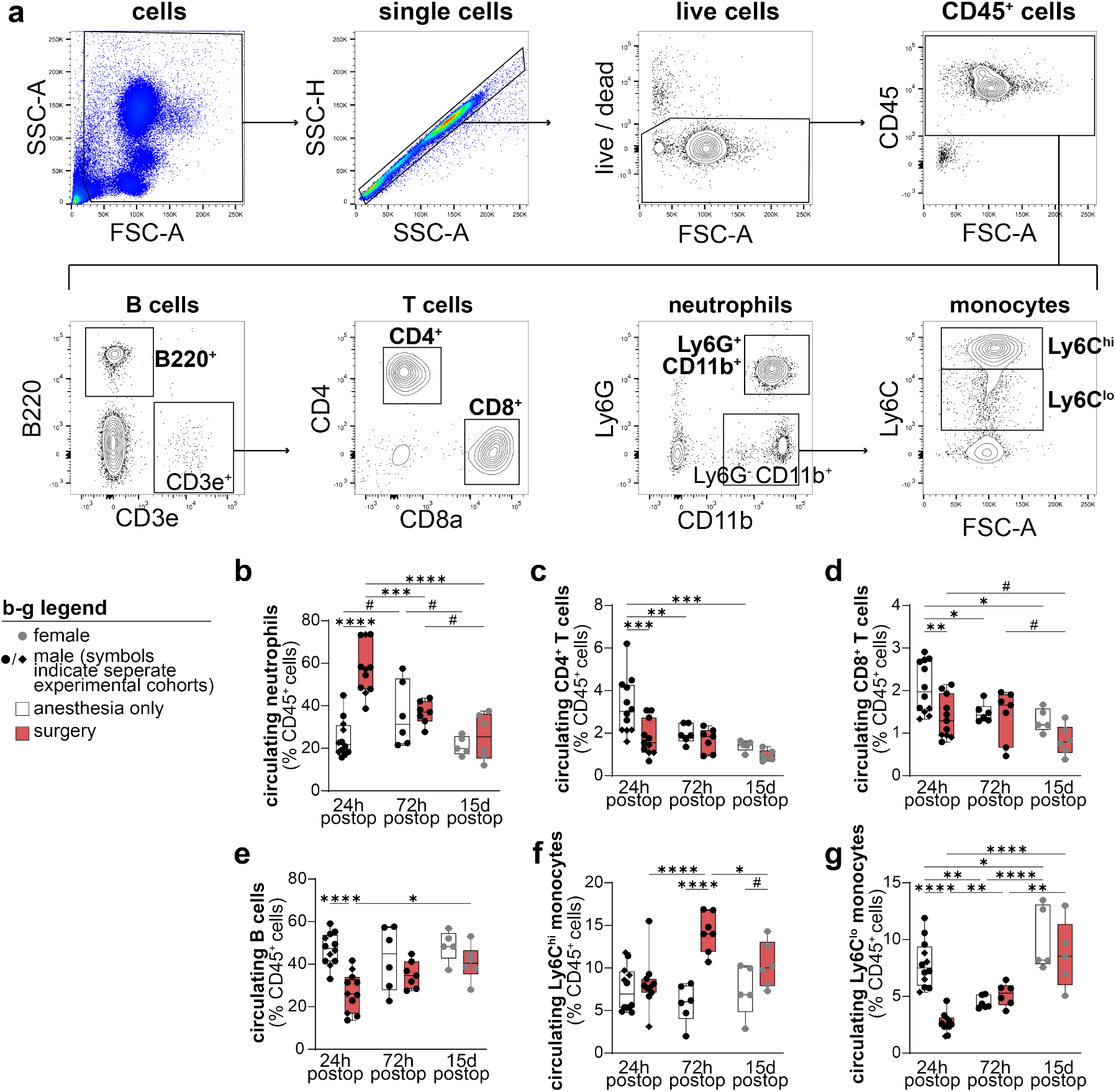
Gating strategy and circulating immune cells. Circulating immune cells from blood obtained by cardiac puncture were gated for flow cytometry as follows (**a**): cells, single cells, live cells, CD45^+^ cells. Subpopulations were gated from CD45^+^ cells as follows: B cells B220^+^; cytotoxic T cells CD3e^+^CD8a^+^; helper T cells CD3e^+^CD4^+^; neutrophils CD11b^+^Ly6G^+^; inflammatory monocytes CD11b^+^Ly6G^-^Ly6C^hi^; patrolling monocytes CD11b^+^Ly6G^-^Ly6C^lo^. Cell populations as percentage of CD45^+^ cells at 24h, 72h and 15d in anesthesia control and surgery groups are represented (**b-g**). Data as min to max boxplot, # P ≤ 0.1, * P ≤ 0.05, ** P ≤ 0.01, *** P ≤ 0.001, **** P ≤ 0.0001.

**Supplemental Fig 2:**
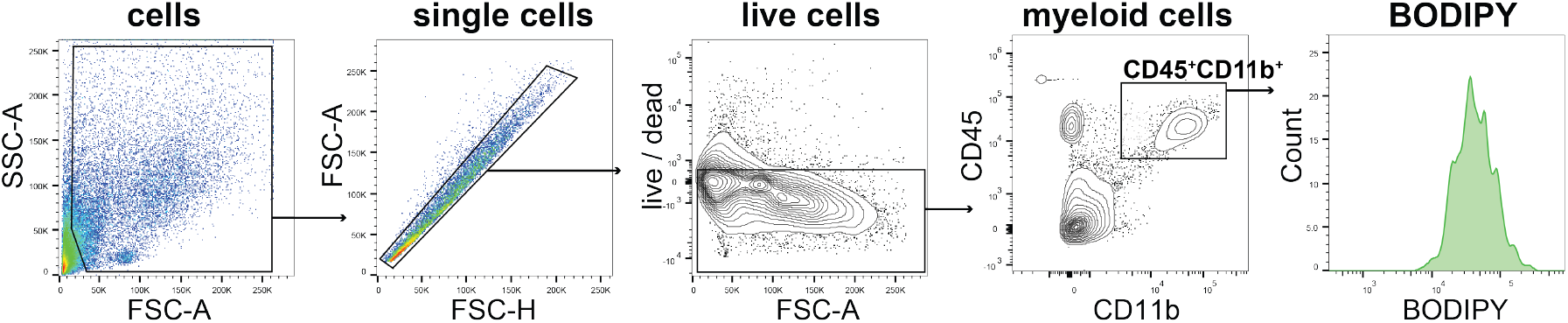
Gating strategy for aortic arches. Cells from aortic digest were gated for flow cytometry as follows: cells, single cells, live cells, myeloid CD45^+^CD11b^+^ cells. Median BODIPY from CD45^+^CD11b^+^ cells was obtained to determine BODIPY MFI.

**Supplemental Fig 3:**
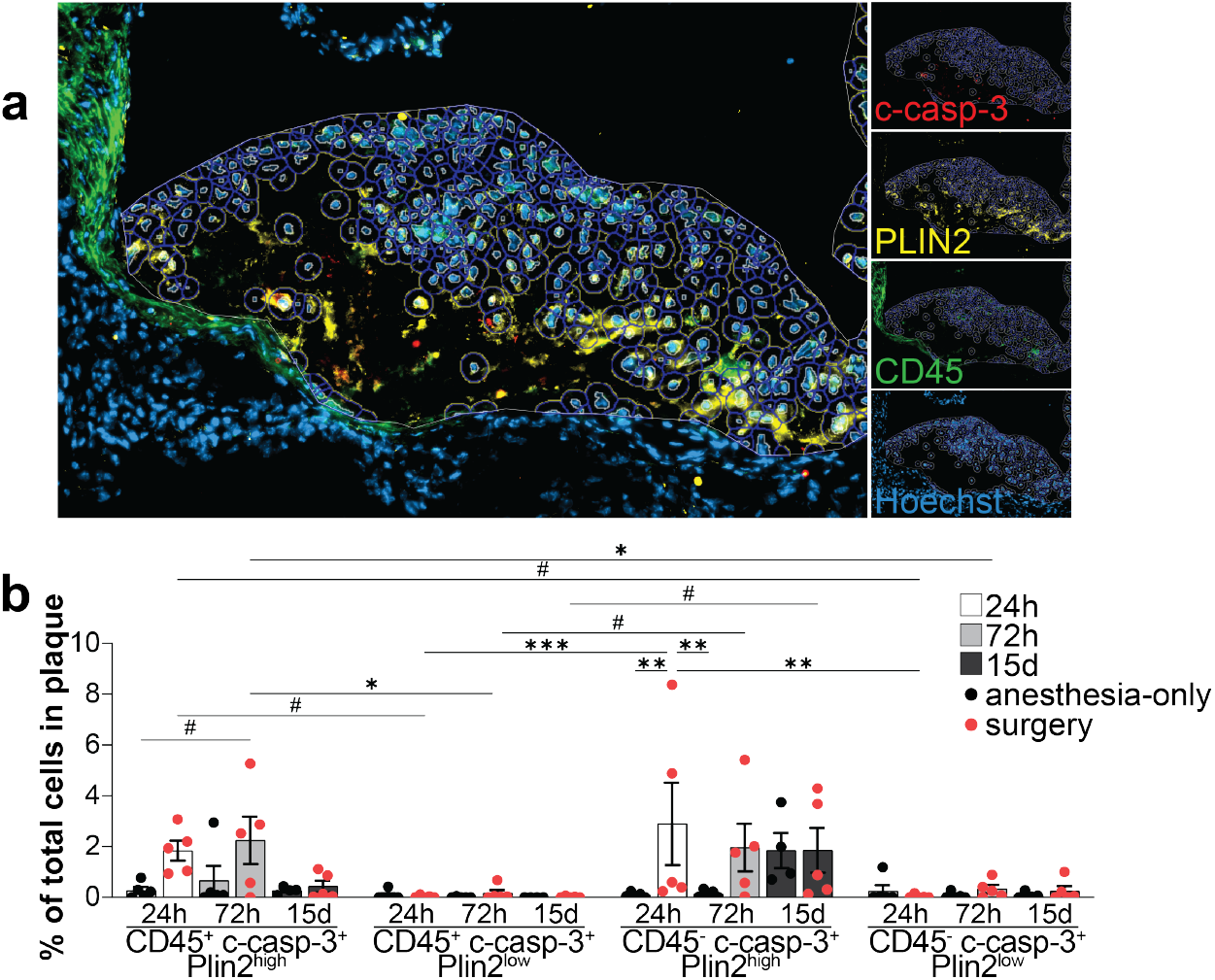
ZenDesk 2D image analysis cell outlines. Fluorescence intensity of each channel was assessed by automated generation of cell outline around identified nuclei, using the ZenDesk 2D image analysis toolkit, as represented (**a**). Expanded data from **Fig.2m**. Apoptotic (c-Casp-3^+^) cell populations defined by expression of CD45 and PLIN2, expressed as percentage of total plaque cells. Data as mean ± SEM, # P ≤ 0.1, * P ≤ 0.05, ** P ≤ 0.01, *** P ≤ 0.001.

**Supplemental Fig 4:**
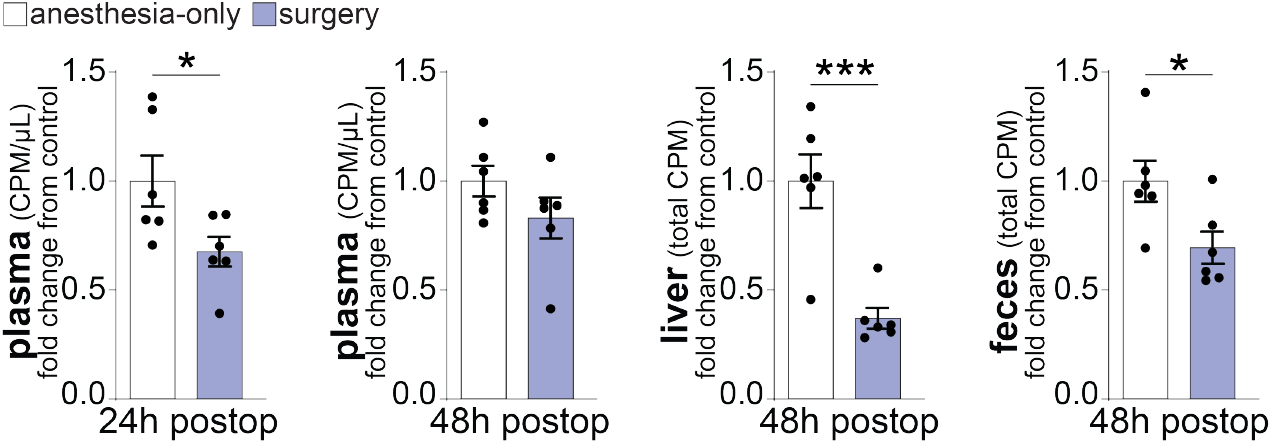
Reverse cholesterol transport in C57BL/6 mice. 12-week-old male C57BL/6 mice (n=6/group) were randomized to anesthesia control or surgery group. BMDMs from sex-& aged-matched C57BL/6 donors were loaded with ^3^H-cholesterol-labeled agLDL for 30h and injected during recovery from anesthesia. Radioactivity was measured in plasma at 24h after surgery, and in plasma, liver & feces at 48h postoperative. Data as mean ± SEM, * P ≤ 0.05, *** P ≤ 0.001.

